# Single Cell Transcriptomics-guided Antisense Treatment Improves Endoderm Differentiation of iPSCs

**DOI:** 10.1101/2020.05.13.092593

**Authors:** P. Hor, V. Punj, Z. Borok, A.L. Ryan (Firth), J.K. Ichida

## Abstract

Directed differentiation of induced pluripotent stem cells (iPSCs) enables the production of relevant cell types for studies of human biology, disease-modeling, and efforts towards cellular therapy and transplantation. However, the low yield and purity of desired cell types can limit the utility of this approach. Enhancing differentiation purity can require extensive optimization of morphogen treatments, and this can still be ineffective for iPSC lines with biased or aberrant differentiation propensities. To address these limitations, we have developed a new approach for increasing the purity of iPSC directed differentiation cultures called <u>R</u>NA sequencing and <u>A</u>ntisense-assisted <u>D</u>ifferentiation (RAD). We performed trajectory analysis of single cell RNA sequencing during iPSC differentiation into endoderm to identify transcription factors responsible for committing cells to undesired, non-endodermal fates. Specific suppression of these transcription factors using antisense oligonucleotides (ASOs) increased the percentage of endodermal cells by up to 20-fold in 3 different iPSC lines that exhibited poor endoderm differentiation. Moreover, this approach required minimal culture manipulation. Thus, RAD improves the utility of iPSCs for basic and translational studies.

## Introduction

Although directed differentiation of induced pluripotent stem cells (iPSCs) into somatic tissues holds great promise for translational applications, the low yield of target cell types limits the utility of this approach (Loh et al., 2014; Loh et al., 2016; Veres et al., 2019). A single morphogen can have different effects on one cell state *versus* the next in a developmental trajectory (Loh et al., 2014; Loh et al., 2016). Thus, epigenetic heterogeneity and asynchronous differentiation kinetics between cells can drive differentiation into unintended lineages (Bock et al., 2011; Kim et al., 2011; Loh et al., 2014; Osafune et al., 2008). Precisely-timed, short-duration administration of morphogens can increase differentiation yield by confining treatment to a period in which most cells are in the appropriate state (Loh et al., 2014; Loh et al., 2016). However, this approach requires extensive optimization of the timing of morphogen treatment, can be labor-intensive, and may require considerable adjustment among iPSC lines (Loh et al., 2014; Loh et al., 2016). Moreover, certain iPSC lines possess characteristics that prevent effective optimization of morphogen treatment, such as asynchronous differentiation kinetics between cells or biased differentiation propensities. (Loh et al., 2016; Tsankov et al., 2015). If a more facile method could improve the purity of directed differentiation, particularly for iPSC lines that are difficult to differentiate into the desired lineage, this would increase the scalability and translational utility of iPSC-based approaches.

To this end, we have developed a systematic approach for improving the purity of the target cell type in iPSC directed differentiation called <u>R</u>NA sequencing and <u>A</u>ntisense-assisted <u>D</u>ifferentiation, or RAD. RAD employs RNA sequencing to identify transcription factors that drive unwanted lineages and gene-targeted ASOs to suppress their function. Using endoderm as a target lineage and iPSC lines with a low efficiency of endoderm differentiation, we show that trajectory analysis of single cell RNA sequencing enables the identification of transcription factors that drive unwanted lineages during endoderm differentiation. Administration of antisense oligonucleotides (ASOs) suppresses these transcription factors and significantly improves the purity of the target endoderm population in three different iPSC lines with minimal manipulation. Thus, RAD provides a facile approach for increasing the purity of iPSC directed differentiation that should be adaptable to multiple culture protocols.

## Results

### Identification of contaminating cell types using single-cell RNA sequencing

Previous studies showed that blocking the induction of unwanted lineages during directed differentiation using extrinsic signaling molecules could enhance the production of desired cell types (Chambers et al., 2009; Loh et al., 2014; Loh et al., 2016). However, because iPSCs differentiate asynchronously, the same morphogen can have opposing effects on one cell state *versus* the next cell state in a developmental trajectory (Loh et al., 2014; Loh et al., 2016) and extrinsic signaling molecules can have undesired effects on a subset of cells within the population. We hypothesized that ASO-mediated suppression of key transcriptional regulators of undesired lineages could provide a similar effect without the risk of perturbing an entire signaling cascade inappropriately in some cells.

To test this hypothesis on iPSC-endoderm differentiation, we quantified endoderm production using a human iPSC line that displays a low efficiency of endoderm formation using our previously published protocol (Firth et al., 2014). In this differentiation protocol, definitive endoderm forms at days 4-5 and with increased specification of the foregut endoderm by days 9-11 (Fig. 1A)(Firth et al., 2014). As we have done previously (Firth et al., 2014), we used *FOXA2* and *SOX17* to identify endodermal cells at days 5 and 10.

**Figure 1.**
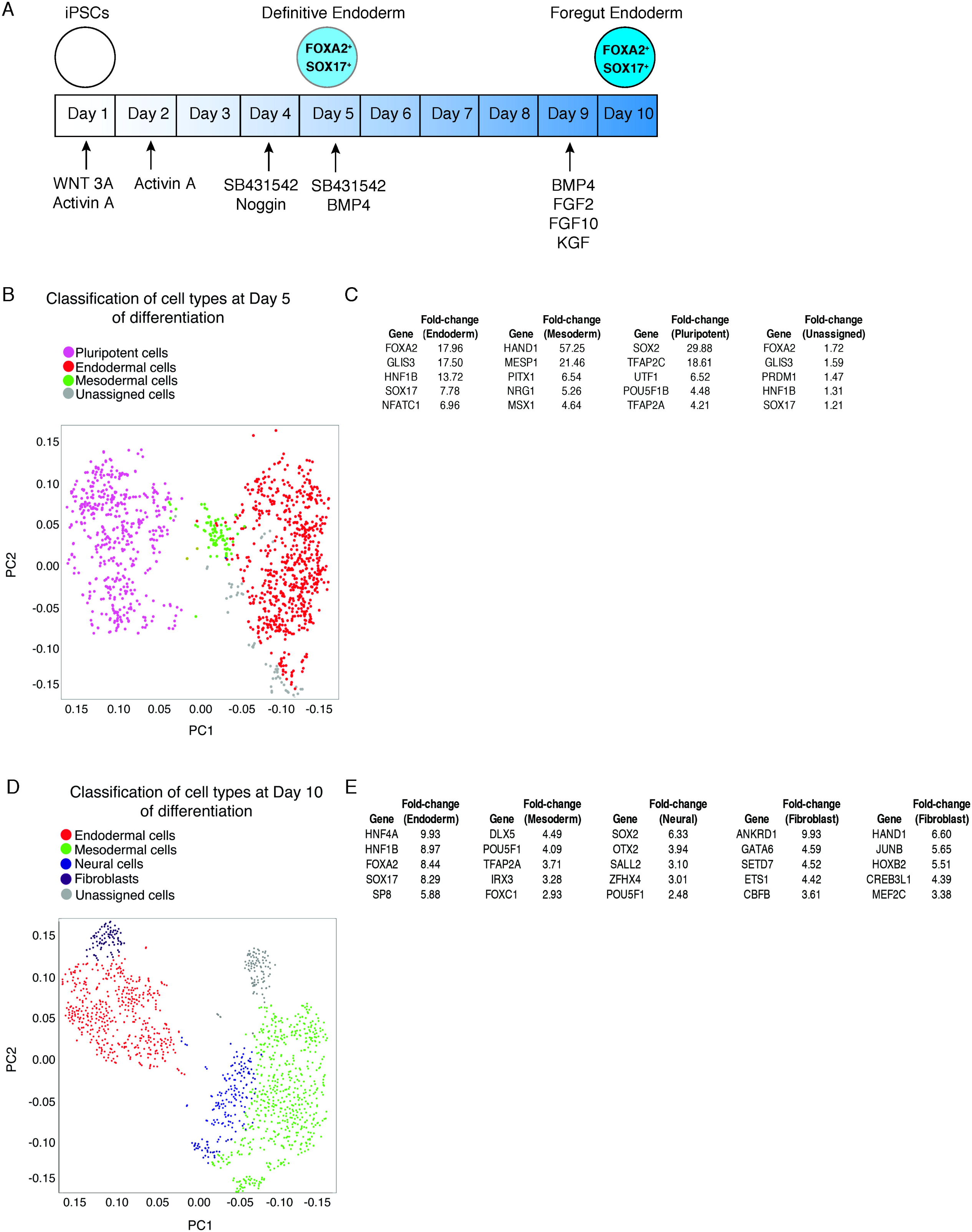
Identification of contaminating cell types in endoderm differentiation. (A) Schematic overview of the iPSC-endoderm differentiation protocol. (B) tSNE plot showing clustering of cells at Day 5 of differentiation. (C) Top-ranked transcription factors by fold-change in each cluster compared to all other cells at day 5 (FDR p<0.05). (D) tSNE plot showing clustering of cells at Day 10 of differentiation. (E) Top-ranked transcription factors by fold-change in each cluster compared to all other cells at day 10 (FDR p<0.05).

To quantify the percentage of endodermal cells at days 5 and 10 of differentiation, we performed single cell RNA-seq analysis of differentiating cultures of iPSC line 1, which we previously derived from fetal foreskin fibroblasts (Firth et al., 2014). 1294 and 1392 cells passed quality control analysis at days 5 and 10 of differentiation, respectively (Fig. 1B, D). Graph-based unsupervised clustering (false discovery rate, FDR p<0.05) distributed the cells into 4 distinct clusters at day 5 and 5 clusters at day 10, suggesting 4 and 5 different cell populations at days 5 and 10, respectively (Fig. 1B-E, Supplementary Tables 1, 2). To classify cell types, we used the unique biomarker genes for each cluster (ANOVA, False Discovery Rate p<0.05, fold-change ± 1.5)(Fig. 1B, D and Supplementary Tables 1, 2). We validated this approach by examining the most differentially-expressed transcription factors in each cluster compared to all other clusters (Fig. 1C, E). At day 5, 45% of cells possessed an endodermal signature, while 43% were endodermal at day 10 (Fig. 1B, D). Thus, fewer than half of all cells adopted the intended endodermal cell identity despite the use of an endoderm-specific differentiation protocol, confirming the low efficiency of endoderm differentiation for this iPSC line.

### Identification of key drivers of contaminating lineages using single cell-RNA seq analysis

We reasoned that eliminating the undesired, non-endodermal lineages would increase the endoderm derivation efficiency. To do so, we designed a workflow using RNA-seq analysis to identify the contaminating lineages and the transcription factors driving them. Because a single morphogen can have opposing effects on one cell state *versus* the next cell state in a developmental trajectory (Loh et al., 2014; Loh et al., 2016), we hypothesized that directly suppressing the transcription factors required for non-endodermal lineages using ASOs might be a more straightforward way of eliminating unwanted cell fates than using extrinsic signaling molecules. We called this approach RNA-seq- and antisense-assisted differentiation, or RAD (Fig. 2A).

**Figure 2.**
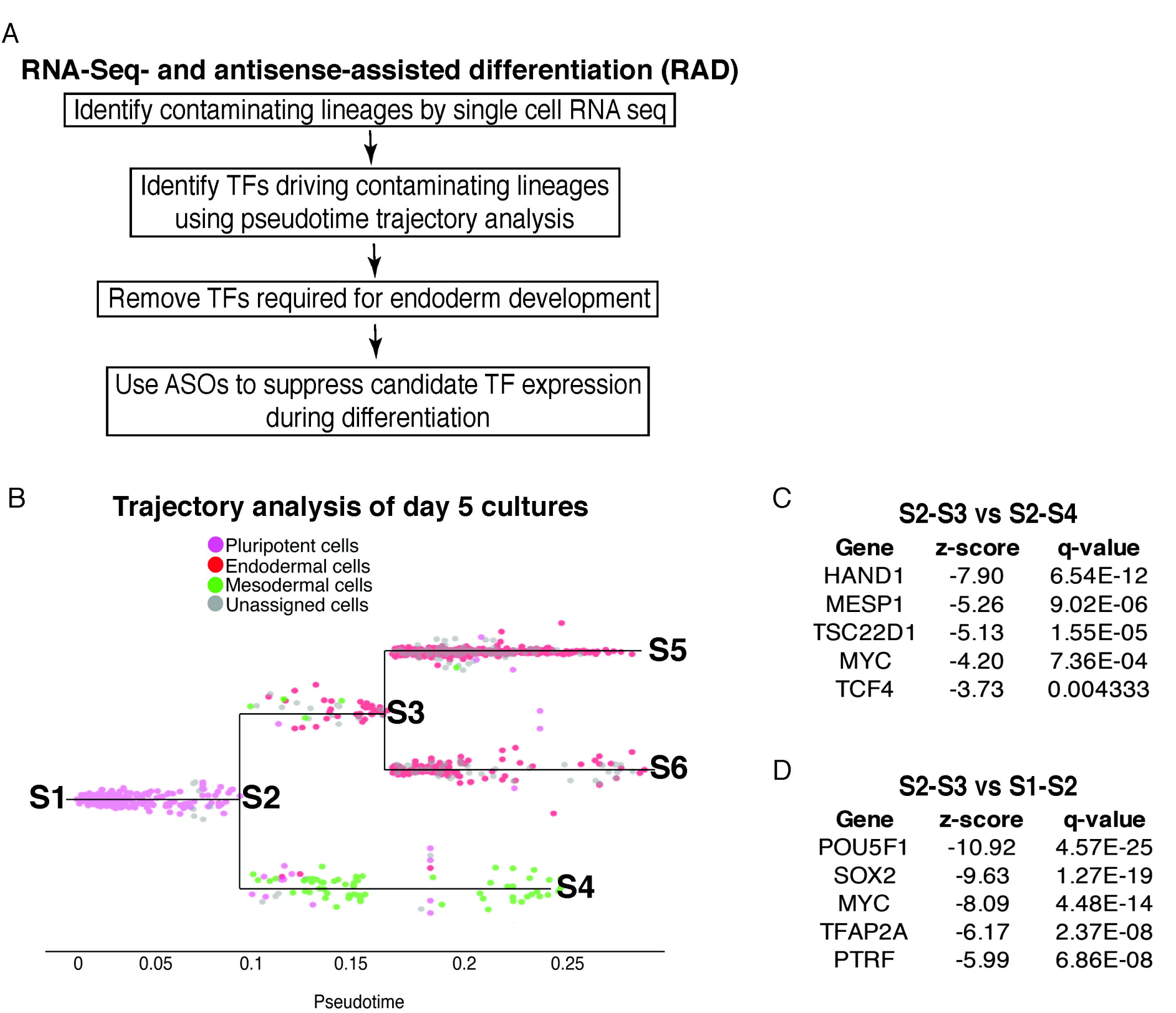
Identification of transcription factors driving contaminating cell lineages. (A) Flow chart outlining the key steps in RAD that together enable the ASO-mediated suppression of contaminating cell lineages during directed differentiation. (B) Pseudotime trajectory analysis based on single-cell RNA sequencing of Day 5 of differentiation. S1-S3 are key branchpoints affecting the production of endoderm. (C) Most differentially-expressed transcription factors by z-score upregulated in S2-S4 (mesoderm) compared to S2-S3 (endoderm). (D) Most differentially-expressed transcription factors by z-score upregulated in S1-S2 (pluripotent) compared to S2-S3 (endoderm).

The single cell RNA-seq data revealed that the largest contaminating populations were comprised of pluripotent cells at day 5 (36.6%) and mesodermal cells at day 10 (35%)(Fig. 1B, D). This was largely due to many cells failing to differentiate or differentiating slowly at day 5, and many cells becoming mesoderm by day 10. Given that the single cell RNA-seq data showed that pluripotent and mesodermal cells comprised the dominant contaminating cell populations (Fig. 1B-E), we focused on suppressing these groups. Since we observed the appearance of mesodermal cells at day 5 (Fig. 1B), we used this time point for further analysis, reasoning that the time point closest to the emergence of mesoderm would be most likely to reveal transcription factors necessary to instruct the formation of mesoderm from mesendodermal cells. Although graph-based unsupervised clustering separated the cells into endodermal, pluripotent, and mesodermal lineages, each population still consisted of cells at different stages of differentiation within a given lineage such as endoderm. Thus, to specifically analyze the transcription factors differentially expressed at critical junctions such as mesendoderm -> mesoderm/endoderm, we performed pseudotime trajectory analysis (Fig. 2B-D).

Pseudotime trajectory analysis comparing the cells at the branch points initiating the endodermal (cells between nodes S2 and S3) and mesodermal lineages (cells between nodes S2 and S4) identified *HAND1* as the most differentially-expressed transcription factor by magnitude of expression difference in the nascent endodermal cells compared to the early mesodermal cells (Fig. 2C, Supplementary Table 3). Comparing the cells at the branch points of endoderm initiation (cells between nodes S2 and S3) and the starting pluripotent state (cells between nodes S1 and S2) showed that *POU5F1* was the most differentially expressed transcription factor in the pluripotent cells compared to the nascent endodermal cells (Fig. 2D, Supplementary Table 4).

Importantly, HAND1 and OCT4 are not required for endoderm development (Matsuura et al., 2006; Sherwood et al., 2009; Tsankov et al., 2015), indicating that their suppression should primarily affect the mesodermal and pluripotent states. Based on our trajectory analysis of day 5 cultures and the notion that HAND1 and OCT4 are dispensable for endoderm differentiation (Fig. 2A-D), we selected *HAND1* and *POU5F1* as our top-ranked candidates for suppression.

### ASO-mediated suppression of *HAND1* and *POU5F1* enhance endoderm differentiation

To determine if suppressing *HAND1* and/or *POU5F1* could increase the purity of endodermal cells during directed differentiation, we designed a strategy to stably reduce their expression using ASOs (Fig. 3A). We designed 20-nucleotide ASOs containing phosphorothioate linkages and 2’-methoxyethyl sugar modifications to enhance their nuclease resistance and binding affinity (Fig. 3B). Using qRT-PCR analysis, we identified two ASOs per gene that significantly reduced expression of *HAND1* and *POU5F1* (Fig. 3C). We previously showed that ASO administration at concentrations of 3-10 μM without transfection reagents can stably suppress gene expression in iPSC-derived cultures for >1 week, and clinical studies show that a single administration of ASOs with similar chemical modifications can suppress gene expression for several months *in vivo* (Becker et al., 2017; Shi et al., 2019; Shi et al., 2018; Wurster and Ludolph, 2018). Because *HAND1* and *POU5F1* are dispensable for endoderm differentiation (Sherwood et al., 2009; Tsankov et al., 2015), we chose to administer the ASOs at the beginning of the differentiation protocol and used either a 10 μM concentration for either *HAND1* and *POU5F1* alone, or a 3 μM concentration when using ASOs targeting both genes simultaneously (Fig. 4A).

**Figure 3.**
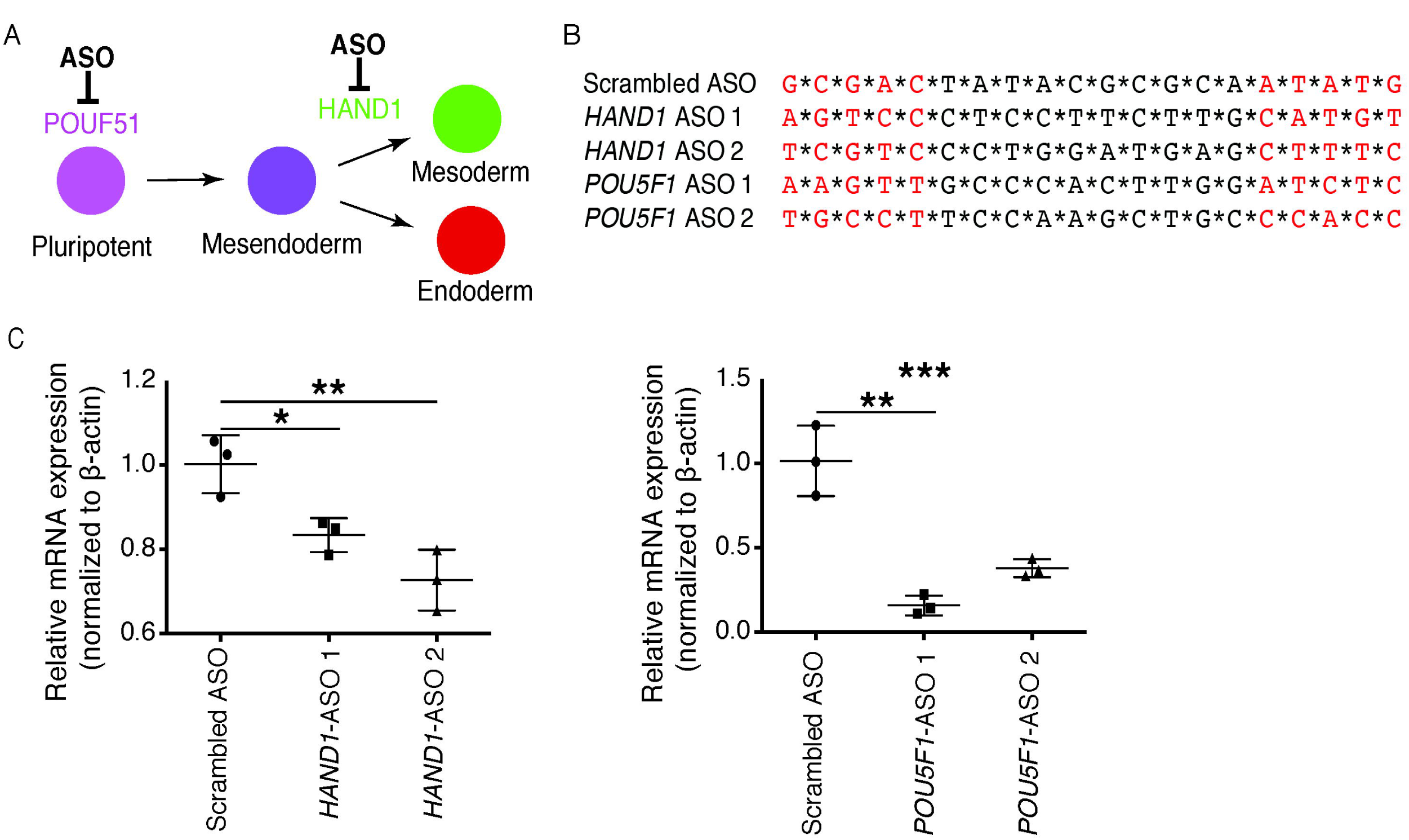
Identification of ASOs that suppress *HAND1* and *POU5F1* expression during endoderm differentiation. (A) Conceptual outline of the ASO-mediated approach to promote endoderm differentiation. (B) ASO sequences. * denote phosphorothioate linkages, red color denotes 2’-methoxyethyl sugar modifications. (C) qRT-PCR analysis iPSC-endoderm differentiation cultures 5 days after two successive treatments (separated by 24 hours) with 3 μM *HAND1* and *POU5F1*. Each data point represents an independent biological replicate. Mean +/- S.D. One-way ANOVA. *p<0.05, **p<0.01. ***p<0.001.

**Figure 4.**
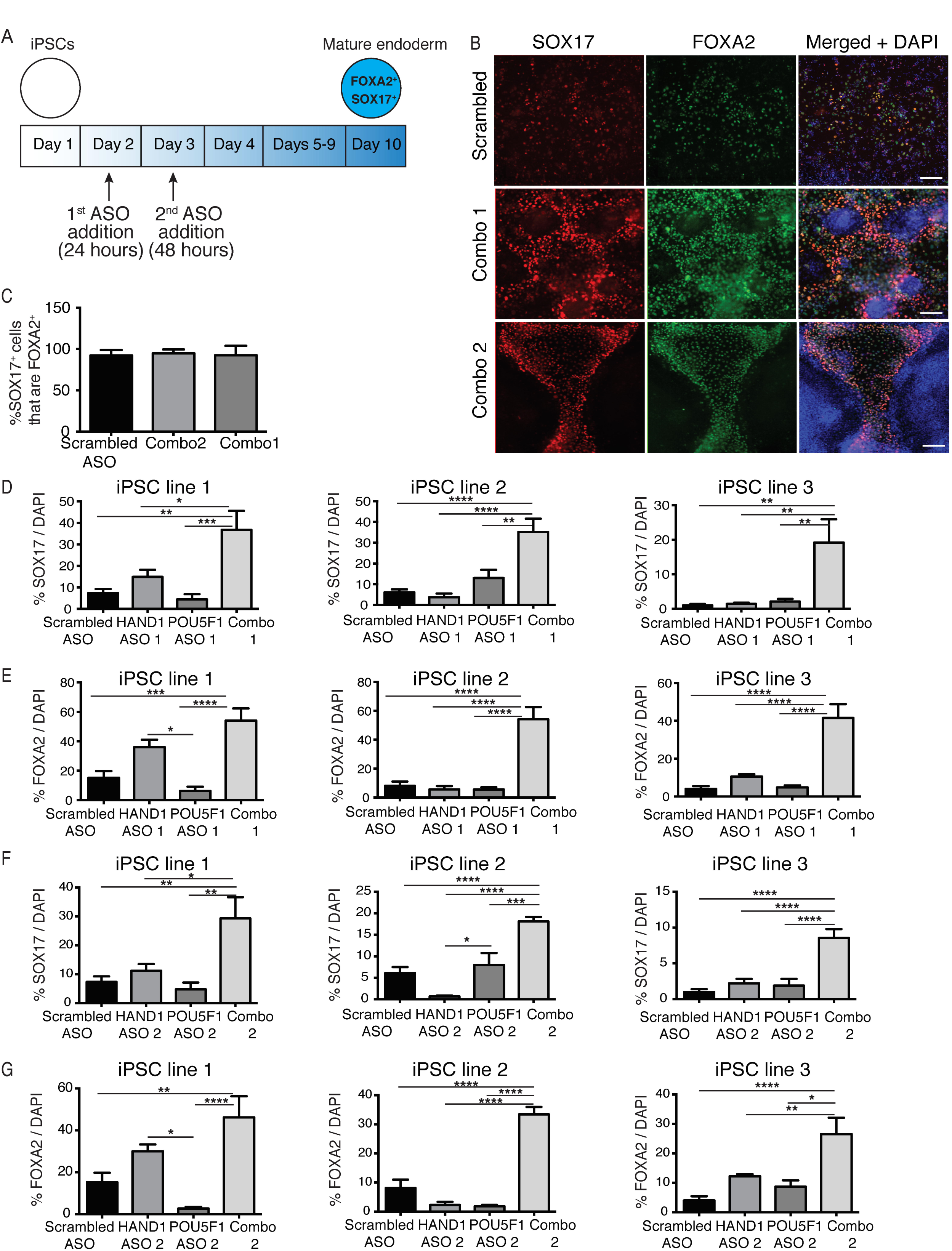
Combined *HAND*1 and *POU5F1* ASO treatment increases the percentage of endodermal cells. (A) Schematic overview of ASO-treatment in the endodermal differentiation protocol. (B) Immunostaining images showing the colocalization of SOX17 and FOXA2 in day 10 endodermal cultures treated with the scrambled ASO, ASO combination 1 (*HAND1* ASO 1 and *POU5F1* ASO 1), or ASO combination 2. Scale bars = 100 μm. (C) Quantification of the immunostaining data showing the percentage of SOX17+ cells that are FOXA2+ after treatment with the scrambled control ASO, ASO combination 1, or ASO combination 2. Mean +/- S.E.M. (D-G) Quantification of SOX17^+^ cells (D, F) or FOXA2^+^ cells (E, G) at day 10 after ASO treatment at days 2 and 3. We quantified 3 independent differentiations for each group for each line. Mean ± SEM, one-way ANOVA. *p<0.05, ** p<0.01, *** p<0.001, **** p<0.0001.

To determine if ASO administration increased the purity of endodermal differentiation, we performed immunostaining for endodermal markers SOX17 and FOXA2 at day 10 (Fig. 4B, C)(Firth et al., 2014). We found that >90% of SOX17+ cells were FOXA2+, and therefore used SOX17 as our primary quantitative readout, with FOXA2 as a secondary readout to validate our findings (Fig. 4B, C). Quantification of immunostaining data showed that the percentage of SOX17+ cells at day 10 in the negative control ASO-treated condition (∼8%) was lower than the percentage of *SOX17*/*FOXA2*-expressing cells we had obtained through single cell RNA-seq analysis (31%, Fig. 1B, D), possibly due to the lower sensitivity of immunostaining analysis (Fig. 4D).

The simultaneous administration of ASOs targeting *HAND1* and *POU5F1* resulted in a 5-fold increase in the number of SOX17+ and FOXA2+ endodermal cells at day 10 in iPSC line 1 (Fig. 4B, D, E). Suppressing only *HAND1* or *POU5F1* alone did not significantly increase the percentage of endodermal cells, indicating that targeting both genes produced an additive or synergistic effect that was required for significant improvement (Fig. 4D, E).

We hypothesized that suppressing *HAND1* and *POU5F1* would improve endoderm differentiation in other iPSC lines that form endoderm with low efficiency. We tested this hypothesis using two additional iPSC lines derived from different donors (iPSC lines 2 and 3)(Fig. 4D-G). ASO treatment targeting *HAND1* and *POU5F1* also increased the percentage of endodermal cells in iPSC lines 2 and 3 by 5- and 20-fold, respectively (Fig. 4D, E). To confirm that these effects were due to suppression of *HAND1* and *POU5F1* and not off-target effects of the ASOs, we repeated these experiments using a different set of ASOs. Again, ASO-mediated suppression of *HAND1* and *POU5F1* significantly improved endoderm differentiation in all 3 iPSC lines (Fig. B, F, G). Thus, simultaneous suppression of *HAND1* and *POU5F1* provides an up to 20-fold improvement in the percentage of endoderm cells and is effective across multiple iPSC lines.

## Discussion

The low purity of target cells after the directed differentiation of iPSCs is a critical barrier to translational applications. Although precisely-timed, short-duration administration of morphogens can improve differentiation purity, this approach requires extensive optimization, may be difficult to scale, and can be ineffective in iPSC lines with low or biased differentiation propensities (Loh et al., 2014; Loh et al., 2016). To help overcome these challenges, we have developed a systematic approach for improving the purity of the target cell type in iPSC directed differentiation called RNA-sequencing- and antisense-assisted differentiation, or RAD. RAD comprises the following key steps: 1) identify the major contaminating cell types using single cell RNA-seq, 2) use pseudotime trajectory analysis to identify transcription factors driving the formation of contaminating cell types, 3) design ASOs to suppress the transcription factors during differentiation. Using RAD, we improved the efficiency of endoderm formation by up to 20-fold and showed that the antisense treatment was effective across multiple iPSC lines that displayed poor endoderm differentiation.

Pseudotime trajectory analysis enabled the identification of transcription factors driving contaminating lineages by highlighting cells at flanking lineage branch points in the differentiation process. Interestingly, we found that pluripotent and mesodermal cells, not ectodermal cells, were the major contaminating cell types in our endoderm differentiation protocol. This suggests that one problem is that many iPSCs exit the pluripotent state more slowly than anticipated and misinterpret or fail to respond to signaling cues administered at days 2-4 of the differentiation protocol. The second problem seems to be that mesendodermal cells differentiate into mesoderm as often as endoderm. This may again relate to the kinetics of differentiation since we observed a wide distribution of cells between the starting pluripotent state and the mesendodermal branchpoint at day 5, suggesting that some cells were not in the appropriate state to interpret the pro-endodermal signaling. Still, we showed that suppressing a single transcription factor per contaminating cell lineage was sufficient to provide a 5-20-fold improvement in endoderm differentiation efficiency.

At day 10, we observed a population of FOXA2+ cells that did not stain positively for SOX17. Other studies have reported similar findings during endoderm differentiation (Wang et al., 2011) and this may reflect asynchronous activation of these markers or different limits of detection by immunostaining. Nevertheless, we observed this FOXA2+/SOX17-cells in both negative control- and *HAND1/POU5F1* ASO-treated cultures, indicating that suppression of *HAND1* and *POU5F1* did not influence this result.

We anticipate that one could easily extend this approach to other target cells or lineages of interest. ASO-mediated suppression of *HAND1* and *POU5F1* successfully enriched cultures for endodermal cells and did not require optimization of treatment duration because it targeted transcription factors that are dispensable for endoderm formation, as opposed to signaling pathways that usually require precisely-timed administration and removal. Thus, RAD provides a facile approach for increasing the purity of directed differentiation cultures that should increase the translational utility of iPSCs.

## Acknowledgments

We acknowledge the use of the services at the USC Optical Imaging Facility, the Genomics Core in the Department of Biomedical Sciences and Dr. Barry Stripp’s laboratory at the Cedars-Sinai Medical Center, the LungMAP consortium, and the high-performance computing cluster at USC. Author contributions: Conceptualization, P.H., V.P., A.L.R, Z.B., J.K.I.; Methodology, P.H., V.P., A.L.R, Z.B., J.K.I.; Investigation, P.H., V.P.; Writing, Review & Editing, P.H., V.P., A.L.R, Z.B, J.K.I.; Bioinformatic analysis, V.P.; Funding Acquisition, J.K.I, Z.B.; Resources, J.K.I., A.L.R and Z.B.; Supervision, J.K.I., A.L.R and Z.B.

## Declaration of interests

J.K.I. is a co-founder of AcuraStem Incorporated. J.K.I. declares that he is bound by confidentiality agreements that prevent him from disclosing details of his financial interests in this work.

All data needed to evaluate the conclusions in the paper are present in the paper and/or the Supplemental Data. Additional data related to this paper may be requested from the authors.

## Methods

### iPSC reprogramming

A human fibroblast-derived iPSC line (iPSC line 1) was obtained as described previously (Firth et al., 2014) and used for all experiments. Two additional human lymphoblastoid cell lines from healthy subjects (ND00184 and ND03719) were obtained from the National Institute of Neurological Disorders and Stroke (NINDS) Biorepository at the Coriell Institute. These lines were reprogrammed into iPSCs as described previously (Okita et al., 2013; Shi et al., 2019; Shi et al., 2018) and labeled as iPSC line 2 and 3, respectively.

### iPSC-endoderm differentiation

On day 0, a single cell suspension of iPSCs was generated using Accutase (Stemcell Technologies, cat. no. 7920) and cells were plated on Geltrex- (Life Technologies, Cat. no. A1413302) coated plates at a density of 1,200,000 cells per well of a 6-well plate. Plated iPSCs were differentiated using a previously published protocol (Firth et al., 2014). On day 1, cells were cultured in RPMI medium (Life Technologies, cat. no. 11875-093) with 25 ng/ml recombinant human WNT3A (R&D Systems, cat. no. 5036-WN-010) and 100 ng/ml Activin A (Humanzyme, cat. no. HZ-1140). On days 2 and 3, cells were cultured in RPMI medium with 1% FBS and 100 ng/ml Activin A. On day 4, cells were cultured in complete serum free differentiation medium (cSFDM, 75% Iscove’s Modified Dulbecco’s Media (IMDM) (Life Technologies, cat. no. 12440053, 25% Ham’s F12 (Life Technologies, cat. no. 11765-054, 0.5X B27 with Retinoic Acid (Life Technologies, cat. no. 17504-044), 0.5X N2 (Life Technologies, cat. no. 17502-048), 0.75% Bovine Serum Albumin (RPI, cat. no. A30075-100.0), 1X Glutamax (Life Technologies, cat. no. 35050). 50 μg/mL ascorbic acid in H2O (Sigma, cat. no. A8960-25G, 4.5×10-4M MTG (Sigma, cat. no. M1753-100ML)) with 500 nM SB431542 (Selleckchem, cat. no. S1067) and 20 ng/ml recombinant human (rh)Noggin (Peprotech, cat. no. 120-10C). On days 5-9, cells were cultured in cSFDM with 500 nM SB431542 and 5 ng/ml Bone Morphogenetic Protein (rhBMP4) (Humanzyme, cat. no. HZ-1045). On days 9-10, cells were cultured in cSFDM with 5 ng/ml rhBMP4, and 10 ng/ml of Fibroblast Growth Factor 2 (rhFGF2) (Peprotech, cat. no. 100-28), Fibroblast Growth Factor 10 (rhFGF10) (Peprotech, cat. no. 100-26), and Keratinocyte Growth Factor (rhKGF) (Peprotech, cat. no. 100-19).

### Single cell RNA sequencing

Day 5 and 10 differentiation cultures were dissociated for 30 mins at 37°C using Accutase to obtain single-cell suspensions. Cells were stained with DAPI and sorted for live single cells followed by cDNA library preparation as per the Chromium Single-Cell 30 Library Kit (10x Genomics) manufacturer’s guide at the Genomics Core of the Department of Biomedical Sciences at Cedars-Sinai Medical Center and sequenced using the 10X Genomics platform on a Next Seq 500. A total of 1502 and 1678 cells were sequenced for Days 5 and 10 respectively at 190-226 million reads per sample. The quality reads were aligned to GRCH38 using STAR aligner. Post normalization expression counts for each gene were collapsed and normalized to unique molecular identifiers (UMI) to generate an expression matrix. Graph based clustering along with t-SNE plots were used to study cell heterogeneity. Biomarker genes of each clusters were identified using an ANOVA test (FDR p<0.05, and fold change ≥1.5). For pathway enrichment analysis, a ranked p-value was computed for each pathway from the Fisher exact test based on the binomial distribution and independence for probability of any gene belonging to any enriched set.

### Pseudo-time trajectory analysis

To identify genes that change as cells undergo transition from one state to another, we reconstructed pseudotime trajectories. R package Slingshot (v 0.3.9) was employed using a log2 normalized count data matrix as described previously (Street et al., 2018). Genes for ordering cells were selected if they were expressed in ≥ 10 cells, their mean expression value was ≥ 0.05 and dispersion empirical value was ≥ 2. To define the developmental trajectories, we used cluster-based MST (minimum spanning tree) analysis to infer the lineage structure. Transcription factors were selected for ASO knockdown based on the magnitude of the expression difference between the desired and undesired lineages at the specified branch point in pseudotime. Transcription factors with the highest expression in the undesired *versus* the desired lineage that were also significantly differentially-expressed were chosen for ASO suppression.

### Antisense oligonucleotide (ASO) treatment

ASO gapmers were designed and produced by Integrated DNA Technologies (IDT) with standard desalting before use. ASOs contained 2′-methoxyethyl sugar modifications and phosphorothioate linkage modifications to increase nuclease stability and retention in cells. ASO gapmers were administered at 3 µM per ASO per treatment whether used alone or in combination. ASOs were first added on Day 1 of differentiation for 24 hrs followed by a media change and second addition on Day 2. ASOs were removed 48 hrs after the second addition.

### qRT-PCR analysis of ASO knockdown efficiency

Total RNA was isolated using the Qiagen RNA Easy Mini Kit (cat. no. 74004, 74134) as per the manufacturer’s protocol. 500 ng of RNA was DNase-treated (Worthington Biochemical Corporation, Lakewood, NJ, cat. no. LS003172) as per the manufacturer’s protocol and reverse-transcribed to cDNA using a Protoscript first strand synthesis kit (New England Biolabs, cat. no. E6560S). Quantitative PCR was performed using SYBR Green (Biorad, cat. no. 1725125) in a 5 µl reaction volume at 50 °C for 2 min, 95 °C for 10 min, 95 °C for 15 s, 60 °C for 1 min for 40 cycles and analyzed using 7900HT SDS software (Biorad). Each run consisted of triplicate technical repeats per experiment. Beta-actin was used as an internal control. The data were normalized to the scrambled ASO-treated condition to quantify the knockdown of *HAND1* and *POU5F1* mRNA expression levels at Day 5 of differentiation after 5 days of antisense oligonucleotide (ASO) treatment. Data are expressed as normalized cycle threshold (Ct) ± SEM. Primers used for qPCR were the following: *β-ACTIN* Forward -5’-AACTCAAGAAGGCGGATGGC-3’, *β-ACTIN* Reverse -5’-CTCCTTAATGTCACGCACGAT-3’, *HAND1* Forward -5’-AACTCAAGAAGGCGGATGGC-3’, *HAND1* Reverse -5’-GGAGGAAAACCTTCGTGCTG -3’, *POU5F1* Forward -5’-CAAAGCAGAAACCCTCGTGC-3’, *POU5F1* Reverse -5’-TGATCTGCTGCAGTGTGGG-3’.

### Immunocytochemistry

Cells were fixed in 3.2% (vol/vol) paraformaldehyde (PFA) for 1 hour at 4°C, washed 3X with phosphate-buffered saline (PBS), and permeabilized in 0.5% Triton X-100 for 1 hour at 4°C. Cells were blocked for 1 hour with commercial CAS block (Thermo Fisher Scientific, cat. no. 008120) and then incubated in primary antibody (Supplementary Table 5) at 4°C overnight. Cells were washed 3X in 0.1% Triton X-100 in PBS and incubated for 1 hour in secondary antibodies. Stained cells were then incubated in DAPI for 5 min before washing and mounting on glass microscope slides in Immunomount (Thermo Fisher Scientific, cat. no. 9990402) or Vectashield with DAPI (Vector Laboratories, Burlingame, CA, cat. no. H-1500). Images were captured on a Zeiss LSM 800 confocal microscope at the USC Optical Imaging Facility (Los Angeles, CA).

### Statistical analysis

All experiments were conducted using at least three technical replicates (e.g., three 6-wells or Transwells) from the same differentiation. All experiments were replicated (independent differentiations) at least three times except where otherwise indicated. Unless otherwise noted, all analyses were performed in GraphPad Prism v.6. All data are presented as mean ± SEM. Statistical significance was determined by one-way analysis of variance (ANOVA) among three or more groups. P < 0.05 was considered statistically significant.

## Supplementary Data

**Supplementary Table 1. Differentially expressed genes in Day 5 cells, 4 clusters, false discovery rate (FDR) p<0.05.** FC = Fold-change.

**Supplementary Table 2. Differentially expressed genes in Day 10 cells, 5 clusters, false discovery rate (FDR) p<0.05.** FC = Fold-change.

**Supplementary Table 3. Differentially expressed genes in Day 5 pseudotime trajectory analysis between cells within branchpoints S2-S3 and branchpoints S2-S4.** The z-score represents the number of standard-deviations that a value is away from the mean expression of all genes in the same group.

**Supplementary Table 4. Differentially expressed genes in Day 5 pseudotime trajectory analysis between cells within branchpoints S2-S3 and branchpoints S1-S2.** The z-score represents the number of standard-deviations that a value is away from the mean expression of all genes in the same group.

**Supplementary Table 5.** Primary antibodies used in immunostaining experiments.

| Name | Brand | Catalog number | Dilutions |
| --- | --- | --- | --- |
| FOXA2 | Cell Signaling Technology | 8186T | 1:500 |
| SOX17 | Thermo Fisher | AF1924-SP | 1:500 |
| FGB | Sigma | HPA001900 | 1:100 |

